# Protein Detection and Localization in Plant Cells Using Spot-Tagging

**DOI:** 10.1101/2020.11.24.396507

**Authors:** Andriani Mentzelopoulou, Chen Liu, Panagiotis Nikolaou Moschou

## Abstract

Fluorescent labelling of proteins without compromising their activity is crucial for determining their spatiotemporal localization while retaining their functionality. Spot-tag is a 12-amino acid peptide recognized by a single-domain nanobody. Here we introduce the spot-tag as a labelling strategy for proteins in fixed and living plant cells, using as an example the microtubule motor centromeric protein E-related Kinesin 7.3. Spot-tagging of ectopically introduced Kinesin 7.3 does not interfere with microtubules and spot staining results in a close-grained fluorophore labelling revealing a localization pattern that resembles “beads-on-a-string”. We anticipate that our protocol will apply to many more demanding protein cellular targets, offsetting activity perturbations and low photon quantum yields imposed by other protein-tagging approaches.

## INTRODUCTION

Most current labelling strategies exploit antibodies or recombinant proteins fused to various stable or photoactivatable fluorescent proteins (FPs) or fluorogenic-labelling enzymes, e.g. the Halo-, CLIP-, or SNAP-tag (Virant et al., 2018). Conventional antibodies introduce significant linkage errors by displacing the fluorophore from the target, while large protein/enzyme tags may affect expression, cellular localization, folding and/or function. Although small peptidic epitopes, e.g FLAG-, HA-, or Myc-tag are widely used, which can be also arranged in tandem arrays to recruit medium-affine binding antibodies, they may not provide sufficient labelling.

Instead of using antibodies, a 15-amino-acid peptide-tag can be probed by high-avid fluorescently labelled monomeric streptavidin, which, however, can be affected by the binding of endogenously biotinylated proteins, especially in plant cells (Arora et al., 2020). Alternatively, reversibly on-/off-binding labels in point accumulation for imaging of nanoscale topography (PAINT) microscopy allow for a continuous and therefore ultra-high-density readout as they are not limited by a predefined fluorophore tagging pattern (Schnitzbauer et al., 2017). Yet, this approach can only be used for structures like membranes or DNA combined with illumination-confined arrangements, e.g. surface-near or light-sheet illuminations.

As a promising substitute for conventional antibodies, small-sized nanobodies (antibody fragments derived from heavy-chain-only camelid antibodies) coupled with organic dyes, e.g. ATTO probes, were recently introduced (Virant et al., 2018). Despite their capability to directly probe endogenous antigens, the *de novo* production of nanobodies and their validation is cumbersome and time-consuming; only a very limited number of microscopy-compatible nanobodies are available by now. Due to their applicability for nanoscopy of widely used FP-fusions, GFP-, and RFP-nanobodies became extremely popular tools. This strategy, however, relies on the correct expression of FP-fusions and does not cope with problems arising from mislocalization, steric hindrances or dysfunction. Thus, nanobodies directed against short and functionally inert tags might prove advantageous.

On par with these approaches, the versatile labelling and detection strategy comprised the short and inert “spot” peptide-tag (PDRKAAVSHWQQ) and a corresponding high-affinity bivalent nanobody has been successfully used in animal cells (Virant et al., 2018). Here we refine the spot-tagging approach for high-resolution imaging of plant cells. We demonstrate the benefits of our approach for protein labelling with minimal linkage errors in fixed and living cells using the paradigmatic component of the centromeric protein E-related Kinesin-Separase Complex (KISC) Kinesin 7.3 (Kin7.3).

## Results and Discussion

### Spot-tag method establishment

Kin7.3 is a core component of the KISC with predominant microtubule (MT) localization. KISC modulates cell polarity and MT dynamics during plant growth. Kin7.3 is partially redundant with Kin7.1 and Kin7.5, which also comprise KISC (Moschou et al., 2016). Firstly, we examined the applicability of the spot tagging method by generating stable *A. thaliana* transgenic lines expressing Kin7.3 fused with His/FLAG epitopes, along with N-terminal spot and C-terminal myc sequences under the meristematic-specific RPS5a promoter (pRPS5a::HF-spot-Kin7.3-myc). Loss-of-function *kin7.1kin7.3kin7.5* (*k135*) mutant leaves are elongated and curly (Supplemental file, Fig. S1). We could see partial complementation in plants expressing pRPS5a::HF-spot-Kin7.3-myc in the *k135* mutant background (Supplemental file, Fig. S1). As live imaging detection of spot fluorescence was only partially successful showing signal only in the outermost root cell layers (Supplemental file, Fig. S2), we assumed that spot cannot be delivered successfully to inner cell layers. We thus developed a protocol for spot-tagging of *A. thaliana* root tip fixed-cells (Fig. 1A). We incubated fixed-roots in fluorescent anti-spot nanobodies for 1 h or overnight and we counterstained spot-labelled roots with anti-myc (Fig. 1B). Using this approach, we could detect colocalized myc/spot signal, but the intensity for both signals was rather low (Fig. 1B), consistent with the reported low native expression of Kin7.3 protein (Moschou et al., 2016). Taken together, these data suggest that spot-tagging is functional in, at least, fixed *Arabidopsis* root cells.

**Figure 1.**
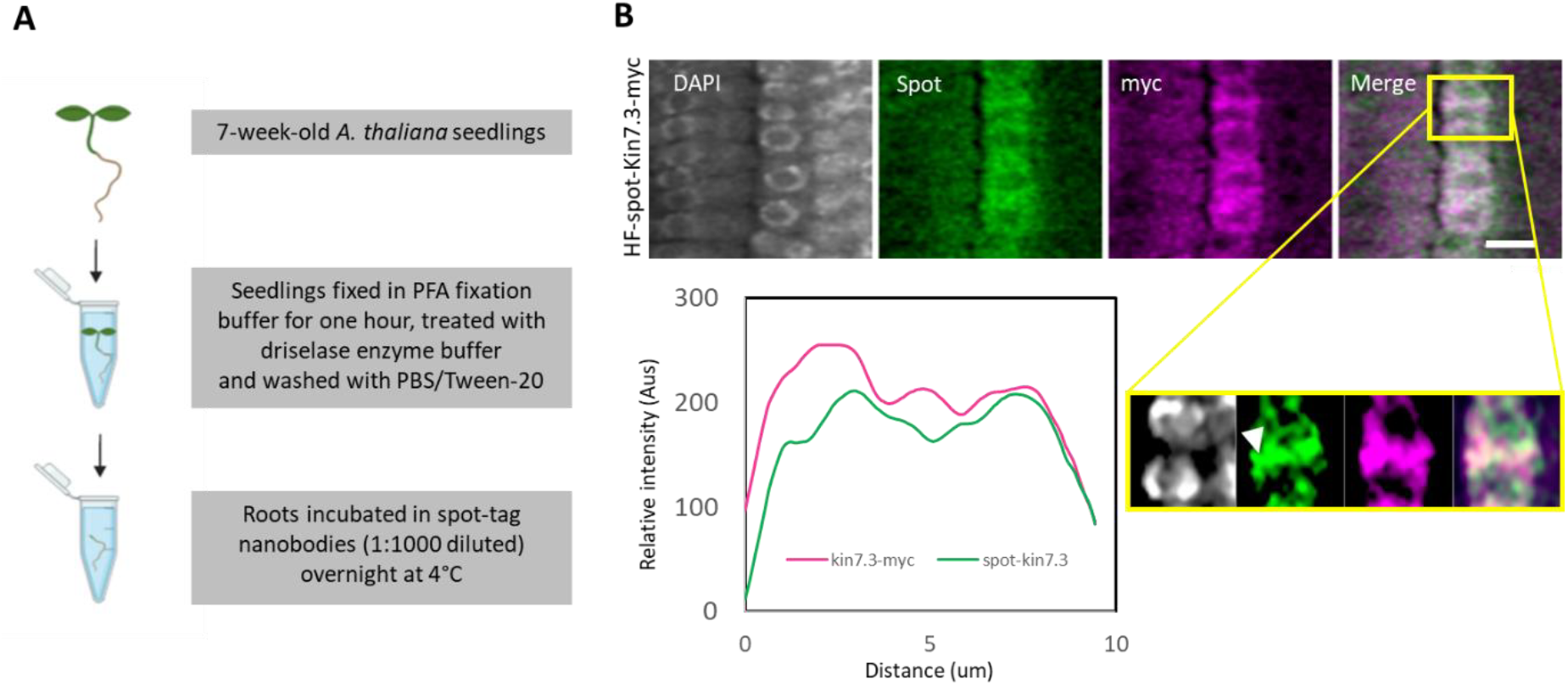
Spot-tag experimental pipeline in stable *A. thaliana* transgenic lines carrying pRPS5a::HF-spot-Kin7.3-myc. **A.** Experimental pipeline for spot detection in fixed root meristematic cells. Scale, 10 um. **B.** Detection of spot-tag and myc/Rhodamine signal in root epidermis meristematic cells and colocalization analysis using plot profile. Pixel colocalization analysis using Pearson correlation coefficient was ~+0.5.

### Kin7.3 detection by spot nanobodies in *Nicotiana benthamiana* leaves

We strived to detect Kin7.3 using spot in the transient mesophyll protein expression system of *N. benthamiana*. We first attempted infiltration of the spot-tag in intact tissue of *N. benthamiana* leaves, which had been infiltrated with a construct carrying pro35::spot-Kin7.3. Under this experimental setting, we could not detect the spot signal. Thus, spot nanobodies cannot be taken up by the mesophyll cells, at least in our settings.

Next, to overcome the physical barrier of the cell wall in spot-tag uptake, we introduced small wounds to the leaf tissue. To this end, we used a razor blade and re-incubated the leaves using spot-tag for 1 h. Under this setting, we could detect high-intensity signal on MTs in cells which were closest to the wounds (Fig. 2A). Our results suggest that this simple wounding approach is sufficient to allow the penetration of spot nanobodies into plant cells.

**Figure 2.**
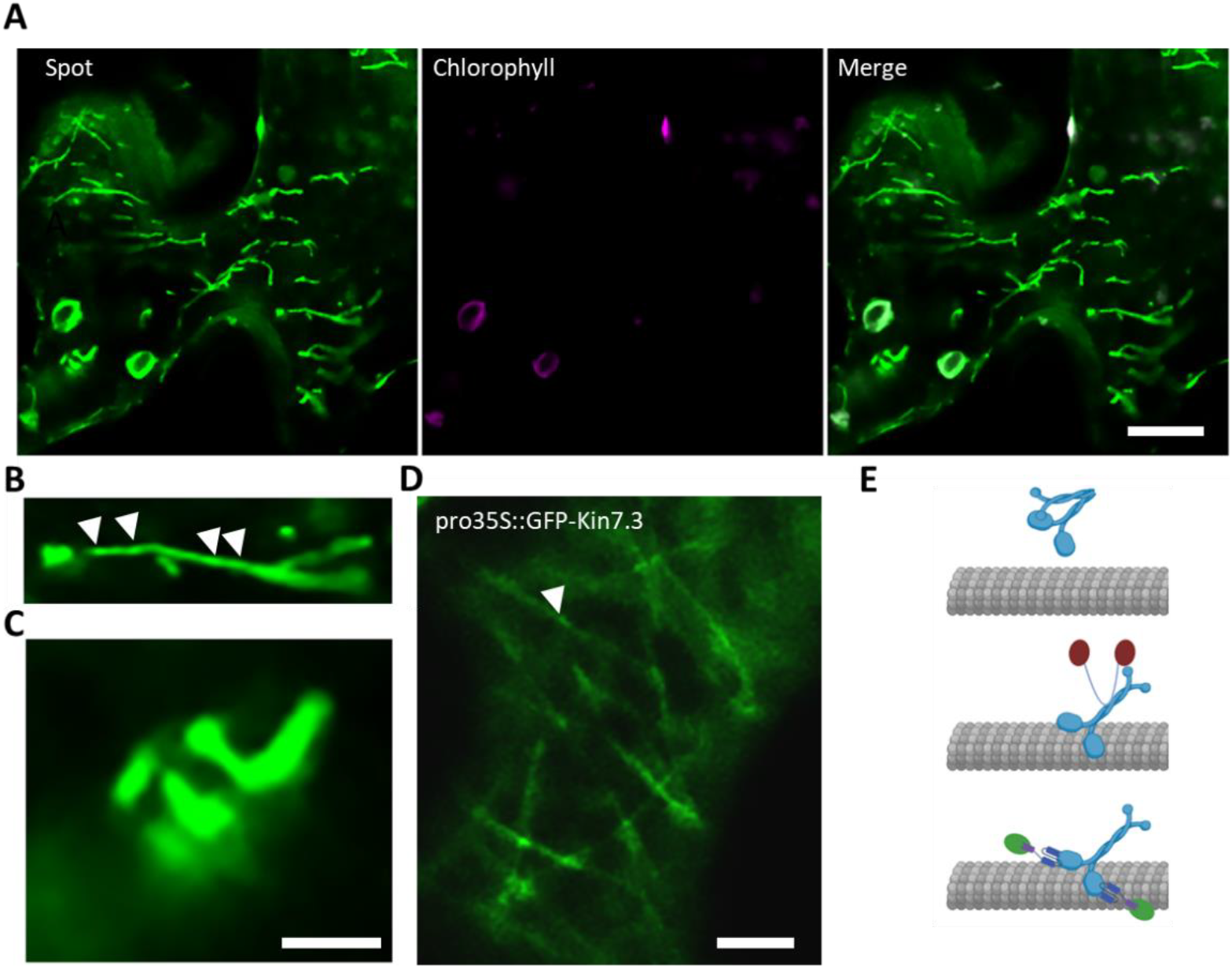
Transient expression and detection of spot-kin7.3 in *N. benthamiana* mesophyll cells. **A.** Spot-tag detection of spot-Kin7.3 expressed in wounded but alive *N. benthamiana* mesophyll cells. **B.** “Beads-on-a-string” detection of spot-Kin7.3 signal (denoted with white arrowheads). **C.** Spiral arrangement of MTs imposed by tension due to overstabilization effect and/or overaccumulation of spot-Kin7.3. **D.** Detection of GFP-Kin7.3 expressed under 35S promoter. Note the increased cytoplasmic signal. **E.** Model of Kin7.3 activation and binding of MTs. In the absence of Kin7.3 or spot nanobodies, Kin7.3 is in the closed conformation (upper). Separase finds on the non-motor C-terminus tail domain, thereby blocking N/C-termini interaction (middle). Spot nanobodies bind on the N-terminus of spot-Kin7.3 leading to a sterical hindrance that allows Kin7.3 binding on MTs.

### Kin7.3 shows a ‘‘beads on a string’’ localization atop cortical MTs

We carried out a thorough examination of MT-based localization of Kin7.3-spot signal. Subcellular localization of Kin7.3 is not affected by the spot-tagging as almost all the signal resided on MTs (Fig. 2A). Surprisingly, Kin7.3-spot signal decorated unevenly the MTs (Fig. 2B), in a pattern reminiscent of “beads-on-a-string” localization observed for other MT-or actin-related proteins (e.g. (Buschmann et al., 2015, Wang et al., 2019)) or proteins that mediate ER-plasma membrane contacts (Wang et al., 2014). On the contrary, using the pro35S::GFP-Kin7.3 under the same conditions (Fig. 2C, D), we could see the increased signal of GFP-Kin7.3 in the cytoplasm suggesting that GFP leads to stereochemical hindrance restricting localization of Kin7.3 on MTs. Moreover, some MTs were under tension perhaps due to the MT overstabilization effect of Kin7.3, leading to a spiral-like MT arrangement (Fig. 2C). Furthermore, as reported previously, the MTs that were decorated by Kin7.3-spot were attached to the plasma membrane, suggesting that Kin7.3 may be involved in plasma membrane-MTs links. Although efficient MT-binding by Kin7.3 depends on the open conformation of Kin7.3 were N- and C-termini of Kin7.3 are far apart due to C-terminal binding of separase, the spot-nanobody may function in the same way by blocking the intramolecular N/C-interaction of Kin7.3 allowing increased MT-binding (model in Fig. 2E).

## Conclusion

We foresee that our spot-tagging approach of can be widely applicable in plant biology, while the two available conjugated to spot fluorophores (i.e. ATTO488/561) will allow detection in at least two different emission wavelengths easing colocalization studies. The high quantum yield of spot-conjugated ATTO probes and their photostability may allow gaining detailed and high-resolution images of spot-tagged proteins, for which fluorescent-images cannot be easily obtained due to low expression levels and/or photostability or quantum yields. Hence, the spot-tagging approach may enable single-molecule localization microscopy (SMLM) techniques, providing outstanding spatial resolutions surpassing limitations, such as poor photon emission or detection efficiency, low fluorophore labelling densities, linkage errors or steric hindrances.

## Materials and Methods

### Plant material and growth conditions

*Arabidopsis thaliana* Col-0 plants were grown on vertical square plates containing half-strength Murashige and Skoog medium (Duchefa), supplemented with 1% (w/v) sucrose and 0.7% (w/v) plant agar. *Nicotiana benthamiana* plants were, directly, grown on soil. Plant growth was done at 22°C on a 16-hr/8-hr light/dark cycle and light intensity of 150 μE m^−2^s^−1^ in a photostable growth chamber (FitoClima 6001.200; Aralab). *Arabidopsis* plants were transformed according to Clough and Bent (Clough & Bent, 1998) using *Agrobacterium tumefaciens* strain GV3101 carrying pRPS5a::HF-spot-Kin7.3-myc construct in a pGWB-based vector (Nakagawa et al., 2007).

### Molecular Biology and Vectors Construction

Kin7.3 coding sequence was amplified from *Arabidopsis thaliana* first-strand cDNA template via RT-PCR. The PCR amplification product was cloned in pDONR221 and subsequently in destination Gateway vectors by BP and LR Gateway recombination reactions, respectively. Sequences of the primers used in this study are available in Supplemental Table 1. The spot tag sequence introduced was PDRVRAVSHWSS.

### Agroinfiltration

*Agrobacterium tumefaciens* GV3101 strain, carrying pro35S::Kin7.3-spot were cultured at 28°C in Yeast Extract Peptone (YEP) liquid medium (1% (w/v) yeast extract, 1% (w/v) peptone, 0.5% (w/v) NaCl) supplemented with 50 ug/ml Spectinomycin and 50 ug/ml Rifampicin. Bacteria were harvested by centrifugation at 2,800 g for 10 min and the pellet was re-suspended in buffer containing 10mM MES pH 5.7, 10mM MgCl_2_ and 200uM acetosyringone; the bacteria were left agitating for 2-4 hours at 28°C. Agrobacteria were harvested, again, respectively, and infiltrated into mesophyll leaves of *N. benthamiana* at a final OD_600_ of 0.4, while co-expressed with p19 silencing suppressor. Approximately six-week-old *N. benthamiana* leaves were used for the infiltration assay.

### Spot tag detection

Roots from *A. thaliana* pRPS5a::HF-spot-Kin7.3-myc transgenic lines were fixed in fixation buffer for 1 h (3.7% paraformaldehyde (PFA; dissolved in warm ddH_2_O supplemented with 2 drops of 0.1 N KOH), 50mM piperazine-N,N′-bis (2-ethane sulfonic acid) (PIPES), 5 mM Ethylene glycol tetraacetic acid (EGTA), 2 mM MgSO_4_, 0.4% (v/v) Triton X-100). Roots were separated from the seedlings using a razor blade and washed with Phosphate-buffered saline/0.01% (v/v) Tween-20 (PBST) buffer and next, they were treated with driselase enzyme buffer for 7 min (2% (w/v) Driselase (Sigma), 0.4M Mannitol, 5mM EGTA, 15mM 2-(N-morpholino) ethane sulfonic acid (MES), pH 5.6, 1 mM phenylmethylsulfonyl fluoride protease inhibitor (PMSF; Sigma)). Epitope blocking was done for 30 min at RT using 3% (w/v) BSA/PBST. Then, roots were incubated overnight at 4°C in anti-myc-mouse primary antibody (1:250, Roche in blocking solution), washed 3 times in PBST. The roots were incubated overnight at 4°C with the secondary anti-mouse (1:500; Jackson immunoreactive, conjugated to Rhodamine Red) and eba488-10 Spot Label ATTO488 (1:1000, Chromotek) diluted in PBS supplemented with 2% (w/v) BSA. After washing in PBST buffer supplemented with DAPI (4′,6-diamidino-2-phenylindole), roots were mounted in Vectashield (Vector Laboratories) mounting medium.

*N. benthamiana* leaves were harvested 3 days post infiltration with the corresponding Agrobacteria carrying the constructs of interest (i.e. spot-Kin7.3 or GFP-Kin7.3). Small cuts from the infiltrated regions were incubated for one hour in anti-Spot-Tag VHH/Nanobodies conjugated to the organic fluorophore ATTO488 (Chromotek), diluted 1:1000 in 10 mM Tris-HCl solution pH 7.5, adjusted with HCl.

The fluorescent images (micrographs) were acquired at RT using the Leica SP8 inverted confocal microscope, with 40x objective (N.A.=1.2), pinhole adjusted to 1 airy unit, and mounting medium ddH_2_O. The excitation wavelength was 488 nm (Emission 506nm - 544nm) for the spot tag detection and 552 nm (Emission 562nm - 636nm) for anti-myc/Rhodamine detection and 405nm (Emission 412nm - 467nm) for DAPI (nuclei detection). The excitation wavelength for chlorophyll was 488 nm (Emission 764nm - 769nm). For fixed-cell imaging, we used 40x oil corrected objective (N.A.=1.3; reflective index n20/D 1.516).

### Image analysis

The image and pixel analyses were done using IMAGEJ software (https://imagej.nih.gov/ij/). Plot profile was calculated along an interactively applied line, and data of intensity measurements were exported to Microsoft EXCEL (Microsoft, Redmond, WA, USA) and plotted. Default modules and options were used. Images were prepared using Adobe PHOTOSHOP (Adobe, San Jose, CA, USA).

## ACKNOWLEDGEMENTS

Part of this work was funded by the start-up grants from IMBB-FORTH.

## REFERENCES

Arora D, Abel NB, Liu C, et al., 2020. Establishment of Proximity-dependent Biotinylation Approaches in Different Plant Model Systems. The Plant Cell, tpc.00235.2020.

Buschmann H, Dols J, Kopischke S, et al., 2015. Arabidopsis KCBP interacts with AIR9 but stays in the cortical division zone throughout mitosis via its MyTH4-FERM domain. Journal of Cell Science 128, 2033–46.

Clough, S. J., & Bent, A. F. (1998). Floral dip: A simplified method for Agrobacterium-mediated transformation of Arabidopsis thaliana. Plant Journal, 16(6), 735–743. https://doi.org/10.1046/j.1365-313X.1998.00343.x

Moschou PN, Gutierrez-Beltran E, Bozhkov PV, Smertenko A, 2016. Separase Promotes Microtubule Polymerization by Activating CENP-E-Related Kinesin Kin7. Dev Cell 37, 350–61.

Nakagawa T, Kurose T, Hino T, et al., 2007. Development of series of gateway binary vectors, pGWBs, for realizing efficient construction of fusion genes for plant transformation. Journal of Bioscience and Bioengineering 104, 34–41.

Schnitzbauer J, Strauss MT, Schlichthaerle T, Schueder F, Jungmann R, 2017. Super-resolution microscopy with DNA-PAINT. Nat Protoc 12, 1198–228.

Virant D, Traenkle B, Maier J, et al., 2018. A peptide tag-specific nanobody enables high-quality labeling for dSTORM imaging. Nature Communications 9, 930.

Wang P, Hawkins TJ, Richardson C, et al., 2014. The plant cytoskeleton, NET3C, and VAP27 mediate the link between the plasma membrane and endoplasmic reticulum. Curr Biol 24, 1397–405.

Wang P, Pleskot R, Zang J, et al., 2019. Plant AtEH/Pan1 proteins drive autophagosome formation at ER-PM contact sites with actin and endocytic machinery. Nature Communications 10, 5132.

