## Supplemental File for "Protein Detection and Localization in Plant Cells Using Spot-Tagging"

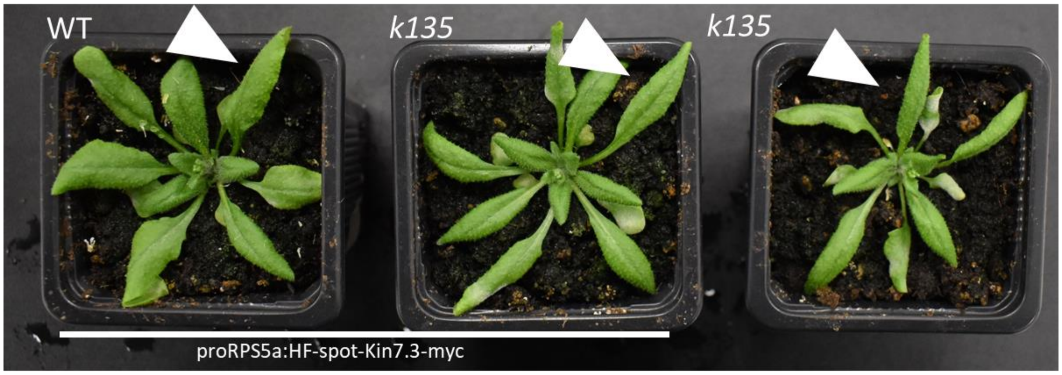


**Figure S1. Complementation of *k135* mutant *Arabidopsis thaliana* line by the pRPS5a::HF-spot-Kin7.3-myc construct.** Arrowheads indicate different degrees of curly leaf-phenotype.

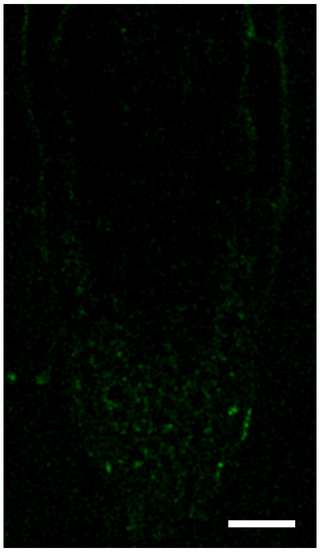

**Figure S2. Live cell imaging of spot-tagging approach and short-term incubation with the spot-tag.** Non-fixed seedlings were incubated in spot nanobodies conjugated to fluorescent dye ATTO488 for 1 h. Representative image of an experiment replicated twice (scale bar 10um).

**Table S1. Primers used for spot-kin7.3 BP cloning.**

| Primer name | Sequence |
|---|---|
| **Spot-kin7.3-F** | AAAAGCAGGCTTAATGCCTGATAGAGTTAGAGCTGTTTCTCATTGGTCTTCTGG ATCTATGGCATCTAGACAAGGAT |
| **Spot-kin7.3-R** | aagaaagctgggttTAAATGAACCTTCCGTTTGTT |
